## Supplementary data for "Emerging variants of SARS-CoV-2 NSP10 highlight strong functional conservation of its binding to two non-structural proteins, NSP14 and NSP16"

**This PDF file includes:**

Figures S1 to S3
Tables S1 to S3
Legends for Datasets S1 to S2

**Other supporting materials for this manuscript include the following:**

Datasets S1 to S2
PDB validation report for 8BZN

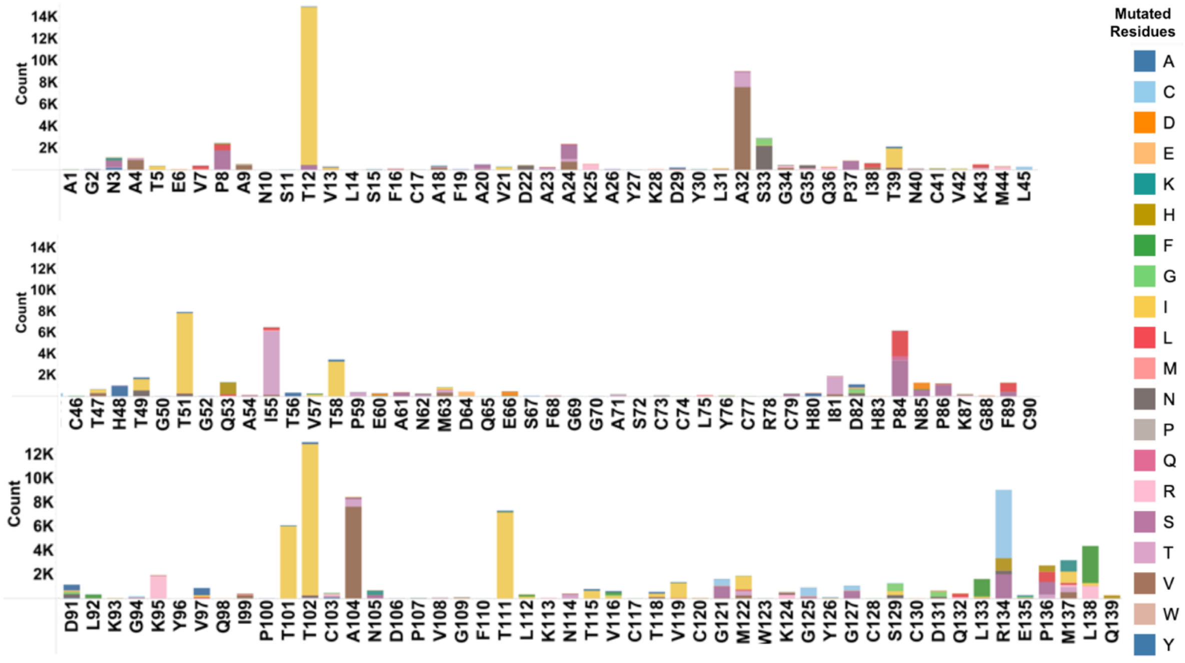


**Fig. S1.** Mutation count of all 820 mutations in NSP10 extracted from 7,070,539 sequences, arranged from residue 1 to 139. The sequences have been saved in Table S1. Different colors represent various mutated amino acids.

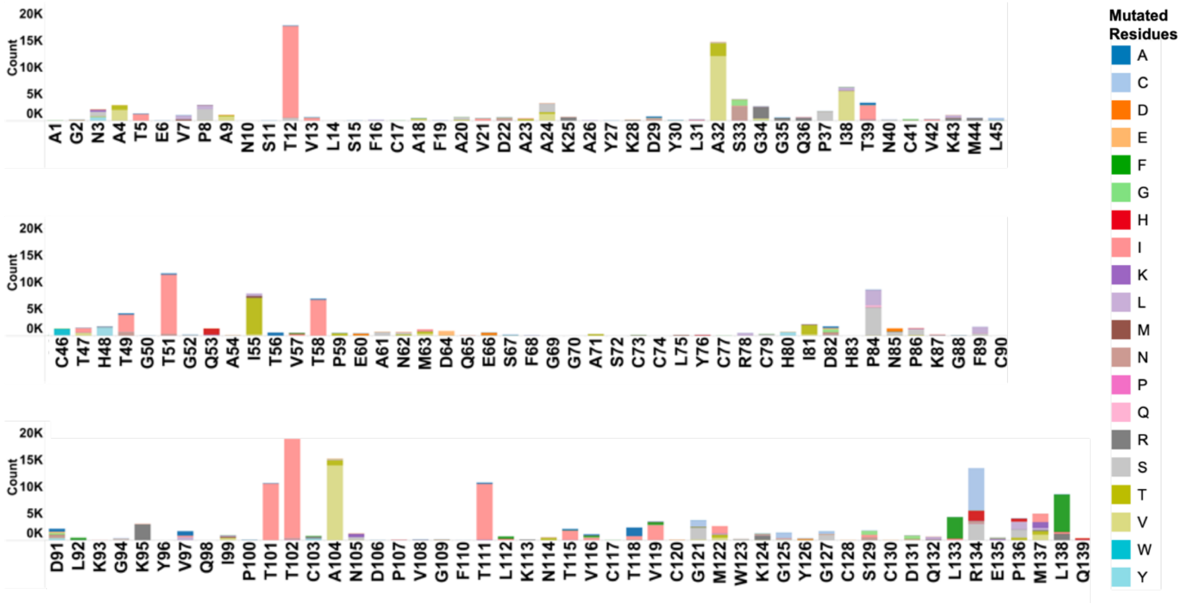


**Fig. S2.** Mutation count of all 878 mutations in NSP10 extracted from 13,032,424 sequences, arranged from residue 1 to 139. The sequences have been saved in Table S2. Different colors represent various mutated amino acids.

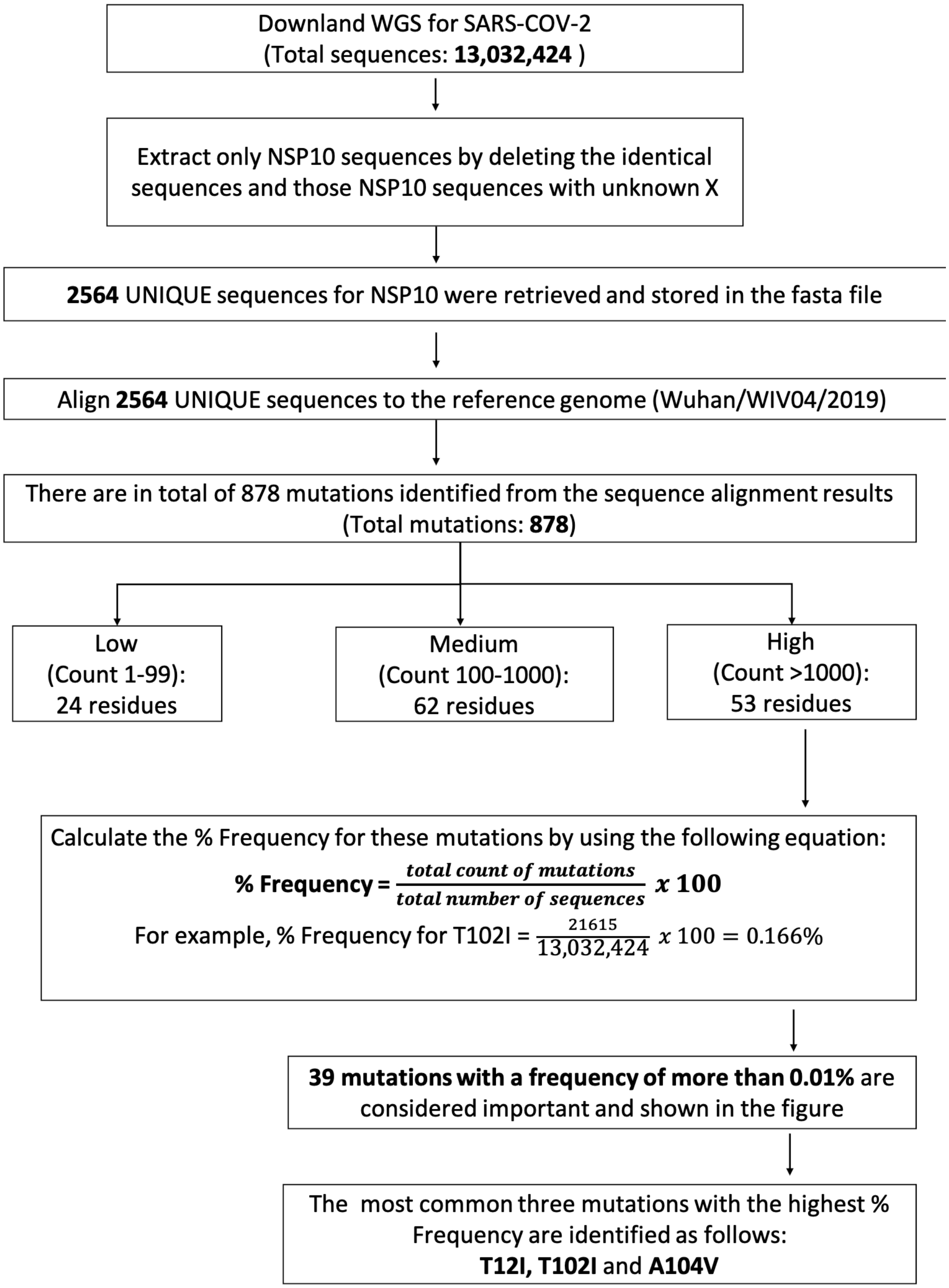


**Fig.S3.** Protocol used for whole genome sequence analysis.

**Table S1:** A detailed table listing all 820 mutations extracted from 7,070,539 sequences. The table enlists the sequence accession no., SARS-CoV-2 strain, position, mutated residue, frequency and % frequency. (Table-S1.xlsx)

**Table S2:** A detailed table listing all 878 mutations extracted from 13,032,424 sequences. The table enlists the sequence accession no., SARS-CoV-2 strain, position, mutated residue, frequency and % frequency. (Table-S2.xlsx)

**Table S3:** Summary of the most frequent variants occurring in NSP10 calculated using Dynamut2. The percent frequency and effect on calculated protein stability are shown. Only mutations with a count larger than 1000 and a percent frequency larger than 0.01% have been included. Residues for which no structure was available has been left blank.

| Mutation | Count | % Frequency | $\Delta\Delta G^{stability}$ (kcal/mol) | Predicted effect on NSP10 stability | NSP10-NSP14 | | NSP10-NSP16 | |
|---|---|---|---|---|---|---|---|---|
| | | | | | Interface | Deleterious | Interface | Deleterious |
| **T102I** | **21615** | **0.166** | **-0.11** | **destabilizing** | **N** | **N** | **N** | **N** |
| **T12I** | **17120** | **0.131** | **-0.68** | **destabilizing** | **Y** | **N** | **N** | **N** |
| **A104V** | **13988** | **0.107** | **-0.1** | **destabilizing** | **N** | **N** | **N** | **N** |
| A32V | 12110 | 0.093 | -0.79 | destabilizing | N | N | N | N |
| T51I | 11050 | 0.085 | -0.61 | destabilizing | N | N | N | N |
| T101I | 10531 | 0.081 | -0.21 | destabilizing | N | N | N | N |
| T111I | 10420 | 0.080 | 0.57 | stabilizing | N | N | N | N |
| I55T | 6814 | 0.052 | -1.75 | destabilizing | N | N | N | N |
| T58I | 6673 | 0.051 | -0.44 | destabilizing | Y | N | N | N |
| I38V | 5610 | 0.043 | -1.54 | destabilizing | N | N | N | N |
| P84S | 5222 | 0.040 | -0.18 | destabilizing | N | N | N | N |
| T49I | 3194 | 0.025 | -0.62 | destabilizing | N | N | N | N |
| K95R | 2979 | 0.023 | -0.41 | destabilizing | Y | N | N | N |
| P84L | 2896 | 0.022 | -0.29 | destabilizing | N | N | N | N |
| V119I | 2805 | 0.022 | -0.14 | destabilizing | N | N | N | N |
| S33N | 2633 | 0.020 | 0.13 | stabilizing | Y | N | N | N |
| T39I | 2116 | 0.020 | -0.24 | destabilizing | N | N | N | N |
| A32T | 2386 | 0.018 | 0.1 | stabilizing | N | N | N | N |
| G121S | 2256 | 0.017 | -0.47 | destabilizing | N | N | N | N |
| G34R | 2188 | 0.017 | -0.33 | destabilizing | Y | N | N | N |
| T115I | 1789 | 0.014 | -0.59 | destabilizing | N | N | N | N |
| I81T | 1773 | 0.014 | -1.22 | destabilizing | Y | N | N | N |
| P37S | 1679 | 0.013 | -1.33 | destabilizing | N | N | N | N |
| T118A | 1625 | 0.012 | -0.3 | destabilizing | N | N | N | N |
| A24S | 1613 | 0.012 | -1.07 | destabilizing | N | N | N | N |
| H48Y | 1553 | 0.012 | -0.22 | destabilizing | Y | N | Y | N |
| M122I | 1382 | 0.011 | -0.2 | destabilizing | N | N | N | N |
| A24V | 1304 | 0.010 | -1.32 | destabilizing | N | N | N | N |
| G121C | 1239 | 0.010 | -0.34 | destabilizing | N | N | N | N |
| R134C | 7953 | 0.061 | | | N | | N | |
| L138F | 7072 | 0.054 | | | N | | N | |
| R134S | 3056 | 0.023 | | | N | | N | |
| P8S | 2050 | 0.016 | | | Y | | N | |
| R134H | 1894 | 0.015 | | | N | | N | |
| P136S | 1573 | 0.012 | | | N | | N | |
| M137I | 1495 | 0.011 | | | N | | N | |
| P136L | 1446 | 0.011 | | | N | | N | |
| A4V | 2055 | 0.016 | | | N | | N | |
| L133F | 4060 | 0.031 | | | N | | N | |
